# Fatty acid synthesis knockdown promotes biofilm wrinkling and inhibits sporulation in *Bacillus subtilis*

**DOI:** 10.1101/2022.05.31.494136

**Authors:** Heidi A. Arjes, Haiwen Gui, Rachel Porter, Esha Atolia, Jason Peters, Carol Gross, Daniel B. Kearns, Kerwyn Casey Huang

## Abstract

Many bacterial species typically live in complex three-dimensional biofilms, yet little is known about systematic changes to gene function between non-biofilm and biofilm lifestyles. Here, we created a CRISPRi library of knockdown strains covering all known essential genes in the biofilm-forming *Bacillus subtilis* strain 3610. We show that gene essentiality is largely conserved between liquid and surface growth and between two media. We developed an image analysis algorithm to quantify biofilm colony wrinkling, which identified strains with high or low levels of wrinkling that were uncorrelated with extracellular matrix gene expression. We also designed a high-throughput screen for sensitive quantification of sporulation efficiency and performed the first screens of sporulation during essential gene knockdown. We found that all basal knockdowns of essential genes were competent for sporulation in a sporulation-inducing medium, but certain strains exhibited reduced sporulation efficiency in LB, a medium with generally lower levels of sporulation. Knockdown of fatty acid synthesis increased wrinkling and inhibited sporulation. These results highlight the importance of essential genes in biofilm structure and sporulation/germination and suggest a previously unappreciated and multifaceted role for fatty acid synthesis in bacterial lifestyles and developmental processes.

**Abstract Importance:** For many bacteria, life typically involves growth in dense, three-dimensional communities called biofilms that contain cells with differentiated roles and are held together by extracellular matrix. To examine how gene function varies between non-biofilm and biofilm growth, we created a comprehensive library of strains using CRISPRi to knockdown expression of each essential gene in the model species *Bacillus subtilis* 3610, which can develop into a wrinkled biofilm structure or a spore capable of surviving harsh environments. This library enabled us to determine when gene essentiality depends on growth conditions. We also developed high-throughput assays and computational algorithms to identify essential genes involved in biofilm wrinkling and sporulation. Knockdown of fatty acid synthesis increased the density of wrinkles, and also inhibited sporulation in a medium with generally lower sporulation levels. These findings indicate that essential processes such as fatty acid synthesis can play important and multifaceted roles in bacterial development.

## Introduction

Many species of bacteria grow in dense, three-dimensional communities held together by extracellular matrix (1). These communities, often called biofilms, represent a form of multicellularity as they typically contain cells with differentiated roles (2, 3). Although biofilm development has recently gained attention as an evolutionary strategy among microbes (3), many aspects of how biofilm development impacts bacterial fitness remain mysterious. The Gram-positive *Bacillus subtilis* has the ability to transition from a motile to a sessile state and has served as a model organism to study both planktonic and biofilm lifestyles (4). Moreover, when *B. subtilis* encounters stressful environments such as nutrient limitation, it can differentiate into a spore that can withstand harsh conditions until the environment is favorable for growth (5).

While *B. subtilis* has been cultivated in laboratories for over a century, most studies have purposefully focused on laboratory strains that are deficient in biofilm formation, and it remains unclear how findings based on these strains translate to wild strains that can form biofilms. For instance, sporulation has been studied extensively, but almost exclusively in laboratory strains using screens to identify non-essential gene disruptions that affect sporulation. Recently, two genome-scale screens of *B. subtilis* laboratory strain 168 non-essential genes revealed an additional 73 (6) and 24 genes (7) linked to sporulation in addition to the >100 genes that had already been identified, suggesting the potential for further discovery involving this developmental process. Moreover, the role of essential genes in sporulation remains uncharacterized.

*B. subtilis* can live as planktonic cultures or as a biofilm (8, 9). Early in biofilm development, cells grow by consuming nutrients from an agar (colony biofilms) or a liquid (pellicles) surface (4). As development progresses, *B. subtilis* colony biofilms adopt a characteristic wrinkling pattern involving buckling of the multilayered structure perpendicular to the surface, correlated with areas in which localized cell death had previous occurred (10). Wrinkle formation is dependent on extracellular matrix production (11) and wrinkles can transport liquid through the biofilm (12). While some of the genetic regulation underlying wrinkling has been elucidated, the roles of essential genes in biofilm wrinkling and whether essential processes mechanistically connect wrinkling to other aspects of development such as sporulation are largely unknown.

In a previous study, we systematically explored the function of essential genes in *B. subtilis* laboratory strain 168 using a CRISPRi gene knockdown library. We showed that dCas9 induction could be used to control expression of RFP in a titratable manner, resulting in ∼30% expression of the target gene at basal levels of dCas9 induction and gradual titration down to essentially 0% expression with full induction, with repression relatively homogenous across a population of cells grown in liquid culture (13). In a separate study, we demonstrated that CRISPRi can effectively knockdown genes in three-dimensional colonies of a GFP-labeled version of the biofilm-forming strain 3610 (14), indicating that CRISPRi is a useful tool for probing essential gene knockdowns both in liquid and on an agar surface. These genetic tools provide the opportunity to broadly and systematically evaluate the role of essential genes in bacterial developmental processes such as non-biofilm and multicellular biofilm colony growth, biofilm wrinkling, and sporulation.

Here, we study a comprehensive library of essential gene depletion strains in *B. subtilis* strain 3610. We demonstrate effective gene knockdown in biofilms over 48 h, the typical time scale of *B. subtilis* biofilm experiments. We find that the subsets of genes essential for growth largely overlap between liquid and colony growth and between two media. We develop high-throughput assays to quantify biofilm wrinkling and sporulation, which revealed that knockdown of fatty acid synthesis genes or gyrase enhances biofilm wrinkling uncorrelated with changes to matrix gene expression and that knockdown of fatty acid synthesis reduces sporulation efficiency. Together, these findings highlight the utility of systems-scale approaches to elucidate the functions of essential genes in planktonic and community behaviors.

## Results

### CRISPRi is an efficient mechanism for gene repression in a biofilm

To investigate the role of essential genes in colony biofilm formation, which is typically monitored over ∼48 h, we first tested whether CRISPRi repression persisted through 48 h of colony growth on agar surfaces. We utilized a *B. subtilis* 3610 strain expressing RFP along with constitutive expression of a guide RNA (sgRNA) targeting the *rfp* gene and xylose-inducible expression of dCas9 (Fig. 1A, Table S1) (14). We grew this RFP-depletion strain on agar plates with the undefined rich medium LB and on plates with the defined, biofilm-promoting medium MSgg, with various concentrations of xylose to cover a range of induction from basal (no xylose) to full knockdown (1% xylose). *rfp* repression was generally titratable with increasing concentrations of xylose, with a relatively uniform RFP signal across the colony at each xylose concentration (Fig. 1B). In colonies on LB, RFP intensity decreased by ∼20-fold with full knockdown (Fig. 1C). The dynamic range was even greater in biofilm colonies grown on MSgg, with a ∼100-fold intensity reduction (Fig. 1C). Thus, we conclude that CRISPRi is a useful tool for inhibiting the expression of essential genes in *B. subtilis* 3610, on both LB and MSgg media.

**Figure 1:**
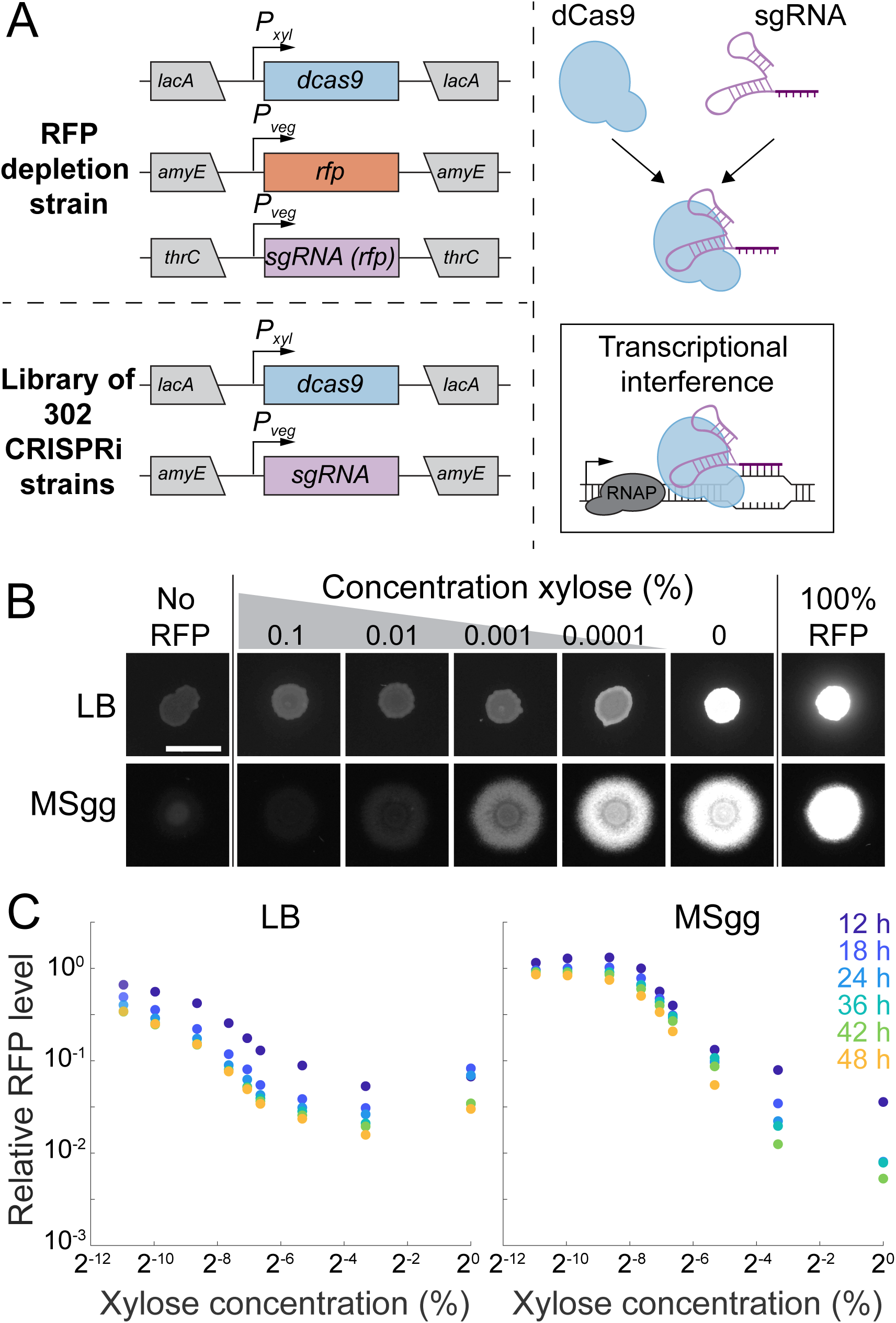
CRISPRi effectively inhibits gene expression during colony growth on LB and MSgg agar. A) In this study, a CRISPRi library was created to individually inhibit the expression of each essential and select non-essential genes. Left: the nuclease-deactivated Cas9 (*dcas9*) gene was inserted under a xylose-inducible promoter (*P_xyl_*) at the *lacA* locus. For the control RFP-depletion strain (top), the *rfp* gene was inserted under the control of a vegetative promoter (*P_veg_*) at the *amyE* locus and a *P_veg_*-*rfp-*targeting sgRNA construct was inserted at the *thrC* locus. For the library of 302 strains, a *P_veg_*- gene targeting sgRNA construct was inserted at the *amyE* locus. Right: dCas9 binds to the sgRNA and inhibits transcription. B) CRISPRi effectively inhibits expression in colonies grown on LB or MSgg and is titratable. Colonies were spotted onto plates with various concentrations of xylose at *t*=0 h and RFP fluorescence at 24 h is shown. All images were contrast-adjusted identically. Scale bar: 5 mm. C) CRISPRi knockdown remains effective for at least 48 h in colonies on LB and MSgg and has greater dynamic range on MSgg. RFP fluorescence was quantified from colonies grown with various concentrations of xylose for 12, 18, 24, 36, 42, and 48 h. RFP levels decreased over time and with xylose concentration.

### Gene essentiality is largely conserved between liquid and colony growth

A previous study determined gene essentiality in *B. subtilis* strain 168 based on the ability to delete the gene of interest during growth in LB (6). Strain 168 has genetic differences relative to strain 3610 in genes important for biofilm formation and other social behaviors such as swarming (15). Nonetheless, we found that the 252 genes described as essential in strain 168 were 100% conserved at the protein sequence level in 3610 (Methods). Thus, we constructed a library of 302 CRISPRi gene knockdowns in *B. subtilis* strain 3610 (Methods, Table S1) (13), which includes the 252 genes identified to be essential in *B. subtilis* 168, 47 conditionally essential and nonessential genes, and 3 controls that do not express an sgRNA. All strains grew as colonies on both LB and MSgg in the absence of inducer (Fig. S2A), demonstrating viability with basal knockdown of any of these genes.

We next determined the extent to which the genes targeted in our library are necessary for growth in our experimental conditions in or on LB and MSgg media (Fig. 2A). In liquid LB+1% xylose to induce dCas9 and fully knockdown the gene target, 111 full knockdowns in strain 3610 did not grow substantially (characterized as OD_600_<0.075 at 5 h) (Fig. 2A, Table S2). In liquid MSgg+1% xylose, 129 full knockdowns in strain 3610 did not grow (characterized as OD_600_<0.075 at 8 h) (Fig. 2A, Table S2), of which 102 also failed to grow in LB+1% xylose (Table S2).

**Figure 2:**
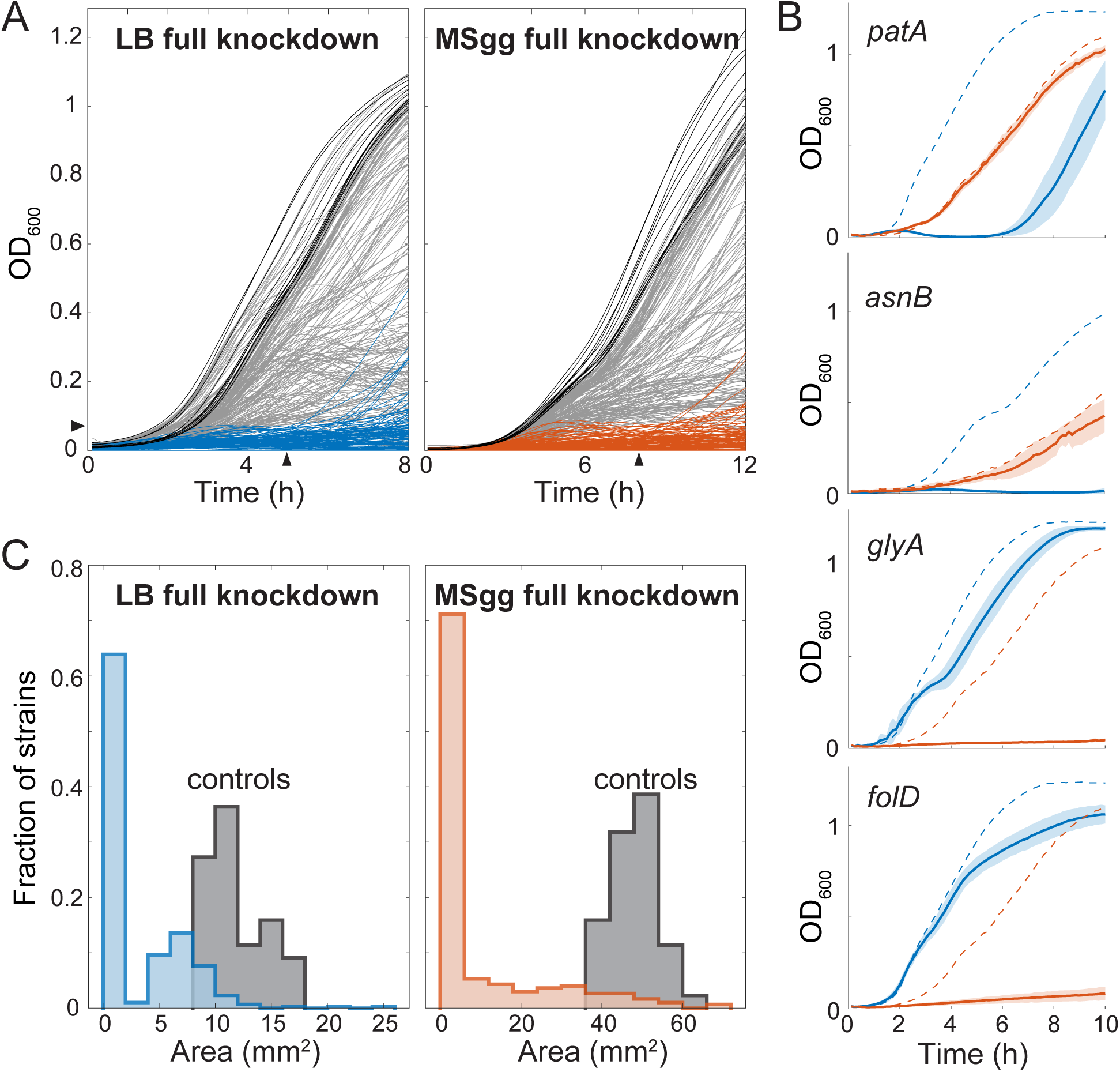
Systematic quantification reveals essential gene knockdowns with media-specific growth phenotypes. A) Full knockdown of many essential genes in liquid media (left, LB+1% xylose; right, MSgg+1% xylose) led to growth impairment. Black lines show the parent control and blue (left) or orange (right) lines show strain 3610 knockdown strains with poor growth (OD_600_<0.075) at 5 h (left) or 8 h (right). B) Four gene knockdowns exhibited dramatically different growth patterns in LB+1% xylose and MSgg+1% xylose. *patA* and *asnB* full knockdowns grew better in MSgg than in LB. *glyA* and *folD* full knockdowns grew better in LB than in MSgg. Dashed lines show the average of the parent control in each medium; the *asnB* full knockdown and the corresponding control were grown at 30 °C, while all other cultures were grown at 37 °C. Blue and orange lines represent growth in LB+1% xylose and MSgg+1% xylose, respectively. Solid lines are the average growth curve over *n*=6 biological replicates, and shading represents 1 standard deviation. C) Full knockdown of many genes resulted in growth impairment on solid media. Plotted are colony sizes at 24 h. Parent controls (*n*=44) are shown in gray.

Under these growth conditions and cutoffs, we identified 27 genes that were potentially required for growth in MSgg but not in LB and 9 genes that were potentially required in LB but not in MSgg. Closer inspection revealed that many of these hits were false positives, as growth of the full knockdown was slightly above the cutoff in one medium; only 4 of the 36 were true positives (Fig. S2B). Full knockdown of *patA* and *asnB,* which are involved in lysine and asparagine synthesis, respectively, inhibited growth in LB+1% xylose but not MSgg+1% xylose (Fig. 2B, S2B), consistent with the medium dependence of the fitness of *patA* and *asnB* knockdowns when grown on plates in competition with wild-type cells (14). Full knockdown of *glyA* and *folD*, which are involved in glycine and folate synthesis, respectively, inhibited growth in MSgg+xylose but not LB+xylose (Fig. 2B, S2B); *glyA* displayed a medium-dependent competitive fitness when grown on plates with wild-type cells and fitness increased when glycine was added to MSgg+1% xylose (14), confirming that *glyA* is conditionally essential and *glyA* knockdowns require glycine for growth. It remains to be determined whether the *folD* medium-specific phenotype is generally due to medium composition or another factor(s).

Since mutant phenotypes can vary between growth in liquid and on solid surfaces (14), we next investigated full knockdown phenotypes on LB and MSgg agar plates. 193 and 195 strains exhibited severe growth defects when grown on LB+1% xylose and MSgg+1% xylose, respectively, with an overlap of 167 strains between these two subsets (Fig. 2C, Table S3). We defined “severe growth defects” as either the absence of growth, generation of suppressors identified as petal-like projections from the original inoculation region (Fig. 2C, e.g., *accD*), or failure to grow beyond the original inoculation region after 24 h (Fig. S2A, Table S3).

In sum, the majority of the genes targeted in the library that were previously identified as essential in *B. subtilis* strain 168 through deletion studies remained important for growth in 3610 in liquid and on solid media.

### Enhancement of wrinkling in fatty acid and gyrase knockdowns is not correlated with expression of matrix-production genes

The defined, biofilm-promoting medium MSgg promoted a broader range of colony size phenotypes compared to LB (Fig. 3A, S2A), similar to our previous study (14). Since growth on MSgg agar promotes biofilm development and wrinkle formation in wild-type *B. subtilis* strain 3610, we used this medium to screen for mutants with aberrant wrinkling patterns. On MSgg, the CRISPRi library exhibited a broad variety of wrinkling patterns across strains, ranging from flat to more wrinkled than wild type. To quantify wrinkling patterns, we developed an image analysis algorithm and an associated wrinkling metric. Wrinkling intensity was quantified as the number of pixels above a threshold after images were background-subtracted, contrast-adjusted, and binarized (Methods, Fig. 3B, S3B). We used this metric to quantify wrinkling across the library at basal knockdown and identified several strains with lower and higher wrinkling than wild type (Fig. 3C, S3A, Table S4).

**Figure 3:**
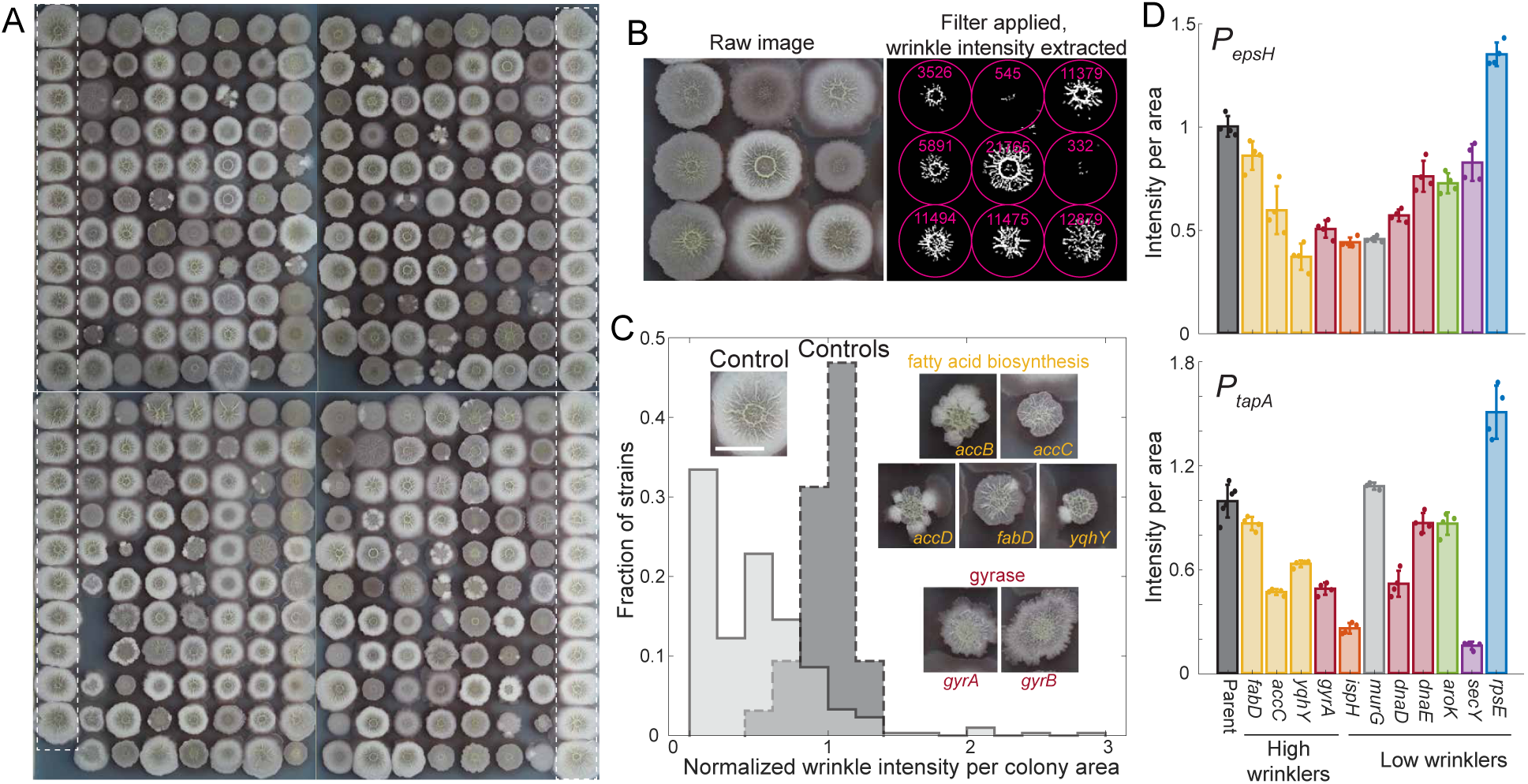
Knockdown of fatty acid synthesis leads to increased wrinkling density. A) Our library of 302 gene knockdowns in strain 3610 covered a variety of phenotypes when grown on MSgg-agar plates for 48 h. Wild-type controls are outlined in white dotted rectangles. The distance between the centers of adjacent colonies is 9 mm. B) Image analysis platform (Methods) automatically quantified degree of wrinkling for each strain. The total wrinkling level (white pixels post-processing) within the colony boundary (pink circle) was extracted for each colony. C) Many strains exhibited significantly lower wrinkling than parent controls, while fatty acid and gyrase knockdowns exhibited higher wrinkling density (wrinkling intensity normalized to colony area). Dark gray dashed histogram represents the parent control, and light gray histogram represents the 302 strains from the gene knockdown library. Insets: representative images of strains with higher wrinkling density; scale bar: 5 mm. D) Matrix gene (*epsH*, *tapA*) expression is uncorrelated with wrinkling density. Reporter levels are shown, normalized so that the average parent control was 1. All *P_epsH_-mNeongreen* gene knockdown strains were significantly different than the parent control (*p*<0.016). Fatty acid mutants are colored in yellow, DNA replication-related mutants are red, ribosome-related mutants are blue, cell wall-related mutants are gray, biosynthesis mutants are green, and a secretion mutant is purple. For *P_tapA_-mNeongreen*, all gene knockdowns except for *murG* were statistically significant (*p*<0.045).

We validated our wrinkling metric by manually curating the positive hits. Nine strains were identified as having high wrinkling intensity per unit area (Fig. S3A). Of these, two (*murAA* and *frr*) turned out to be false positives due to the contrast of the colonies being enhanced by the filter (Fig. S3A). The remaining seven strains clearly exhibited high wrinkling (Fig. 3C, Table S4); of these, *accB, accC, accD,* and *fabD* have defined roles in fatty acid synthesis and the *yqhY* mutant has recently been connected to fatty acid synthesis (16), although its exact role within the pathway remains unclear. *gyrA* and *gyrB* knockdowns also displayed enhanced wrinkling (Fig. 3C). *gyrA* and *gyrB* encode for subunit A and B of DNA gyrase, respectively. Gyrase relaxes positive supercoils and introduces negative supercoils in DNA, and is important for controlling DNA replication initiation and resolving head-on DNA replication-transcription conflicts (17–19); thus, their knockdown likely results in global gene expression changes. We verified the high wrinkling of these seven strains (Fig. S3C) and chose four for further analysis. Since overexpression of matrix components is known to increase wrinkling (20), we quantified the expression of matrix genes in colony biofilms using promoter fusions to *epsH* and *tapA* (*yqxM*). Interestingly, expression of a reporter gene from the *epsH* or *tapA* promoter was significantly reduced relative to the parent control in all gene depletions that displayed high wrinkling (Fig. 3D, S3D). Thus, increased matrix gene expression is not required for the increased wrinkling of these strains.

In addition, several colonies were much less wrinkled than the controls. We manually identified 38 relatively flat colonies by eye; of these, 37 were also identified by our computational analysis with a reasonable cutoff. Strains targeting ribosomal proteins were enriched among the 38 low wrinklers as compared to the entire library (DAVID analysis, *p*=8×10^-5^). We verified the wrinkling pattern of 13 selected low wrinklers (Fig. S3A,C) and chose 11 candidates to explore further (Table S4). Since reduced matrix expression can lead to flat colonies, we tested matrix expression levels in the flat mutants using the transcriptional fusions to *epsH* and *tapA* (Fig. 3D, S3D). Knockdown of *aroK* resulted in significantly reduced expression of *epsH* and *tapA* (Fig. 3D, S3D). By contrast, *rpsE* knockdown colonies displayed higher expression of both matrix genes even though these colonies did not wrinkle (Fig. 3D, S3D), again demonstrating that wrinkling can be decoupled from matrix gene expression in some cases. The remaining candidates displayed a range of expression patterns, most with reduced expression of *epsH* and *tapA* (Fig. 3D, S3D). Thus, our data suggest that flat colony phenotypes may be partly due to reduced matrix levels but that other factors can also play a role.

### A high-throughput assay of sporulation efficiency based on optical density

*B. subtilis* laboratory strains have long been used as a model to study sporulation, and ∼150 non-essential genes required for efficient sporulation and germination have been identified through a variety of screening strategies (21–26). Many of these screens relied on transposon insertion screening (7, 27) or on gene knockout libraries (6) and hence focused only on non-essential genes; thus, the role of essential genes in sporulation remains unknown.

Many genes involved in sporulation were identified based on the inability of non-sporulating mutants to produce a pigmented protein that alters the color of spore-containing colonies (6). However, while strain 168 becomes more translucent when sporulation is blocked, we found that known sporulation mutants in strain 3610 remained opaque (Fig. S4A), rendering color-based screening ineffective. Thus, to investigate the role of essential genes in sporulation of strain 3610, we devised a straightforward, optical density-based, high-throughput screening strategy. We grew strains for 24 h and transferred the cultures to 80 °C for 30 min to heat-kill all vegetative cells (Fig. 4A). After the heat-kill, we diluted cultures into fresh medium so that any viable spores would germinate and grow (Fig. 4A). Notably, our screening strategy by itself does not discriminate between sporulation and germination defects, as both defects would manifest as a growth delay in our assay, although germination defects can be inferred when the sporulation efficiency as measured by CFU counts is maintained.

**Figure 4:**
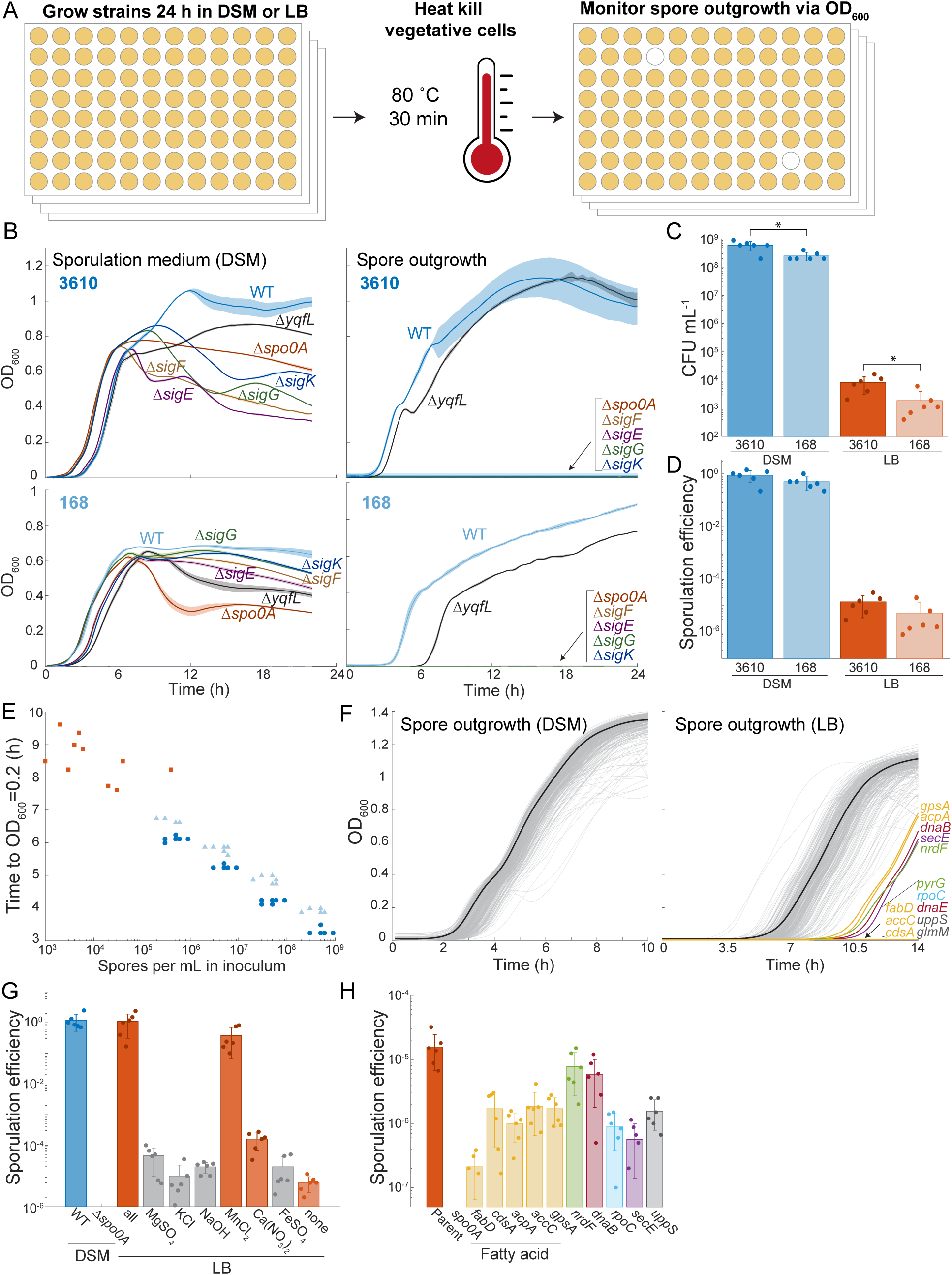
A high-throughput sporulation screen reveals reduced sporulation efficiency due to knockdown of fatty acid synthesis. A) A growth-based, high-throughput assay to identify mutants defective in sporulation. Strains are grown to saturation, vegetative cells are heat-killed, and cultures are inoculated into fresh medium. Any strains without viable spores will not survive the heat kill and hence those wells will remain clear during outgrowth. B) Sporulation mutants exhibit distinct patterns of growth in DSM (left) and are unable to outgrow after heat-killing (right) for both strain 3610 (top) and strain 168 (bottom). Average growth curves over *n*=6 biological replicates are shown as solid lines and shading represents 1 standard deviation. C) There are fewer spores in a 24-h culture of strain 168 compared strain 3610. Cultures were grown in DSM or LB, heat-killed, and plated to measure colony-forming units (CFU). *: *p*<0.02. D) Sporulation efficiency (ratio of CFU post-versus pre-heat-killing) was lower for strain 168 compared with strain 3610. E) Number of spores in the inoculum correlates with growth lag. Cultures post-heat-kill were serially diluted 10-fold and growth curves were measured. The time to reach OD_600_=0.2 was negatively correlated with the number of spores (calculated from CFU). Dark blue circles: strain 3610 grown in DSM, light blue triangles: strain 168 grown in DSM, orange squares: strain 3610 grown in LB. F) All essential gene basal knockdowns are competent for sporulation in DSM, and basal knockdown of fatty acid synthesis reduces sporulation in LB. Spore outgrowth curves following 24 h of growth in DSM (left) or LB (right) and heat-killing. Solid black lines show the average of parent controls (*n*=40 biological replicates), and light gray lines show growth curves of the CRISPRi library. For cultures pre-grown in LB (right), several knockdowns exhibited a delay in growth or did not grow at all (colored lines). Fatty acid mutants are colored in yellow, DNA replication-related mutants are red, ribosome-related mutants are blue, cell wall-related mutants are gray, biosynthesis mutants are green, and a secretion mutant is purple. G) Manganese and calcium nitrate promote sporulation in LB. Sporulation efficiency of strain 3610 in DSM and in LB with various DSM components added. Note that the Δ*spo0A* mutant did not sporulate at all in DSM and hence is not plotted. Sporulation efficiencies in DSM and LB+all additives were statistically indistinguishable (*p*=0.84). Sproulation efficiencies in LB+MnCl_2_ and LB+Ca(NO_3_)_2_ were significantly higher than in LB without any additives (*p*<0.015). H) Fatty acid knockdowns exhibit reduced sporulation efficiency in LB. Colors are the same as in (F). The Δ*spo0A* mutant did not sporulate at all in LB and hence is not plotted. All knockdowns exhibited significantly different sporulation efficiency than the parent control (*p*<0.04).

To validate our assay, we grew known 168 and 3610 sporulation mutants in a sporulation-inducing medium (DSM) for 24 h and monitored OD_600_. Growth curves were reproducible and distinct for the sporulation mutants of each strain relative to wild type, and 3610 reached a higher OD_600_ than 168, likely due to increased oxygen diffusion mediated by surfactin (Fig. 4B, left) (28). Following heat-kill, wild-type cultures displayed robust growth, indicating the presence of spores as expected (Fig. 4B, right). Many sporulation mutants (Δ*spo0A*, Δ*sigF*, Δ*sigE*, Δ*sigG*, Δ*sigK*) displayed no growth after heat-killing, demonstrating the complete absence of spores capable of germination (Fig. 4B, right). Intriguingly, heat-killing of a Δ*yqfL* mutant that is known to undergo reduced sporulation (7) exhibited a substantial increase in lag time after heat-killing relative to wild-type in the strain 168 background, but its lag in the strain 3610 background was similar to wild type, suggesting that this mutant phenotype may be dependent on strain background (Fig. 4B, right). Regardless, these data indicate that our assay successfully identifies mutants fully blocked for sporulation and/or germination and can identify mutants with partially reduced sporulation.

Interestingly, strain 168 exhibited a ∼1 h delay in outgrowth compared to strain 3610 (Fig. 4B, right). After 24 h of growth in DSM, the total number of colony-forming units (CFU) after heat-killing (Fig. 4C) and the sporulation efficiency (ratio of CFU post-heat-kill (spores) relative to pre-heat-kill (spores and viable cells), Fig. 4D) were reduced 2- to 2.5- fold in strain 168 compared to strain 3610. Thus, the reduced number of germination-capable spores produced by strain 168 relative to strain 3610 underlies the increased time until growth was observed in strain 168.

We tested whether outgrowth dynamics after heat-killing could serve as a quantitative proxy for spore counts by measuring the lag time (defined as the time to reach OD_600_=0.2) for 10-fold serial dilutions of strain 168 or strain 3610 after 24 h of growth in DSM and heat-killing. Undiluted inocula exhibited the shortest lag times, and the lag time was highly correlated with spore count (Fig. 4E), with an additional delay of ∼1 h for each 10-fold dilution (Fig. S4B), consistent with a doubling time of ∼20 min. Based on 1 spore being present within the 1 µL inoculum, our assay has a limit of detection of ∼10^3^ CFU/mL of the original culture (Fig. 4E); this limit could presumably be decreased further by increasing the inoculum size. Taken together, these data demonstrate that lowering the number of spores in the inoculum results in predictable delays in outgrowth, and that our assay enables high-throughput quantification of sporulation/germination efficiency.

### All essential gene knockdown strains are competent for sporulation and germination under basal CRISPRi induction in a sporulation medium

To probe whether any strains in our CRISPRi library in the strain 3610 background had reduced ability to sporulate or germinate, we applied our OD-based sporulation assay in basal knockdown conditions after growth in DSM for 24 h. We selected basal knockdown conditions, as full knockdown promotes the emergence of suppressors in strains that are defective for growth (13). During growth in DSM, even though known sporulation mutants displayed distinct growth dynamics (Fig. 4B, right), there was too much variability across the growth curves of the mutants in our library to definitively identify any sporulation mutants based on growth alone (Fig. S4C). Germination dynamics after heat-killing revealed that all strains were competent for sporulation and germination (Fig. 4F, left), indicating that basal expression levels of essential genes are sufficient for sporulation and germination after growth for 24 h in DSM.

### Low sporulation efficiency in LB medium is enhanced through the addition of Mn^2+^ and Ca^2+^

Since all essential gene knockdown strains were capable of sporulation in DSM, we next focused on a medium in which *B. subtilis* shows reduced sporulation efficiency. Strain 3610 exhibited ∼100% sporulation efficiency during growth in DSM. By contrast, in LB, spore CFU/mL and sporulation efficiency of both 3610 and 168 were reduced 10^4^-fold reduced (Fig. 4C,D).

Sporulation requires a combination of starvation and quorum sensing under specific conditions, often involving the undefined medium DSM, which contains Difco nutrient broth and several additional components (Methods). Even though DSM promotes 10^4^-fold more sporulation than LB, growth of wild-type 3610 in DSM resulted in a similar yield as in LB, (Fig. 4C,D, Fig. S4D). A previous study showed that Mn^2+^, one of the components of DSM, is critical for sporulation in a medium to DSM due at least in part to its effects on the enzyme phosphoglycerate phosphomutase (29). To further interrogate the difference in sporulation efficiency between DSM and LB media, we investigated whether the supplements added to DSM affect sporulation in LB. Adding all DSM supplements to LB raised sporulation efficiency to DSM levels, and MnCl_2_ alone was sufficient to raise sporulation efficiency to near DSM levels (Fig. 4G). Furthermore, addition of Ca(NO_3_)_2_ to LB raised sporulation efficiency >10-fold. Thus, robust sporulation heavily depends on the addition of MnCl_2_ and, to a lesser extent, Ca(NO_3_)_2_.

### Lipid metabolism genes are involved in sporulation

We hypothesized that screening in LB, which resulted in vastly reduced sporulation (Fig. 4C,D) due to lower levels of Mn^2+^ compared with DSM, would reveal essential genes with partial sporulation defects. Indeed, screening in LB identified several genes with reduced sporulation efficiency (Fig. 4F, S4E, Table S5). These hits were enriched for genes involved in lipid/fatty acid biosynthesis and metabolism (*fabD*, *cdsA*, *acpA*, *accC*, and *gpsA*; *p*=2.5×10^- 3^, DAVID enrichment score), as well as genes involved in cell wall synthesis (*uppS* and *glmM*), protein secretion (*secE*), transcription (*rpoC*), DNA replication (*dnaB* and *dnaE*), and nucleotide biosynthesis (*nrdF* and *pyrG*). All hits were essential genes except for *gpsA*, which is an auxotroph in strain 168 that is essential in *E. coli* (6) (Fig. 4F, Table S5). Validation of these hits demonstrated that sporulation efficiency was reduced 10- to 100- fold in most cases (Fig. 4H). Intriguingly, despite exhibiting a delay in growth after heat-killing (Fig. 4F, S4E), *nrdF* and *dnaB* had sporulation efficiencies that were only marginally lower than the parent, indicating that the delay may be due to slower germination rather than a reduction in spore number (Fig. 4H). Taken together, screening in a medium with generally lower sporulation efficiency revealed essential genes involved in sporulation and germination.

## Discussion

Here, we used a CRISPRi knockdown library to characterize phenotypes of each essential gene during liquid and colony growth. We found that gene essentiality was consistent between growth in liquid and on a surface as a colony. We also developed high-throughput screening platforms to quantify colony wrinkling and the degree of sporulation in liquid and discovered that fatty acid synthesis was a determinant of the degree of wrinkling and was important for proper sporulation. These findings extend our knowledge of the role played by essential genes during *B. subtilis* growth and development across lifestyles.

Wrinkling is a hallmark property of *B. subtilis* pellicles and colony biofilms. Wrinkles require extracellular matrix, and they are able to transport liquid within the biofilm (12). Despite recognition that wrinkling is a complex feature of biofilms, it is often treated as a binary phenotype for the purpose of classifying mutants. Our image analysis platform enabled quantification of biofilm wrinkling, identifying that knockdown of gyrase or genes related to fatty acid synthesis increases wrinkling per unit area (Fig. 3C). Although wrinkling requires matrix expression and excess matrix can increase wrinkling (30), the knockdowns that we identified increased wrinkling in a manner uncorrelated with matrix expression (Fig. 3D). Thus, the mechanism(s) by which reducing fatty acid synthesis or gyrase activity increases wrinkling remains mysterious and should be the subject of future study.

Sporulation is an important developmental process in a broad range of Gram-positive bacteria that has been extensively studied in *B. subtilis* laboratory strains. More than 150 nonessential genes have been linked to sporulation using screens that relied either on a change in the color of colonies of non-sporulating mutants or, more recently, on transposon sequencing (7). We found that sporulation mutants in strain 3610 remain opaque, motivating the design of a new sporulation screen based on growth following heat-killing that provides a highly quantitative readout of sporulation efficiency (Fig. 4). Using this screen, all essential gene basal knockdowns were sporulation-competent in the sporulation-inducing medium DSM (Fig. 4F); future CRISPRi systems that eliminate (or at least drastically reduce) the emergence of suppressors will enable the interrogation of sporulation during full knockdown of essential genes. In LB medium, sporulation efficiency was generally lower, and fatty acid mutants were among the knockdowns defective for sporulation (Fig. 4F). Notably, our screening strategy does not discriminate between sporulation and germination defects, and two mutants that exhibited near wild-type sporulation efficiency were nevertheless identified by our screen (Fig. 4F,H), likely because they were delayed in germination, highlighting a strength of our assay. Unfortunately, the sporulation efficiency in LB is sufficiently low (∼1 spore in 10^5^ cells) that a manual search for spores to track their germination dynamics is prohibitive without a way to enhance the fraction of spores in the starting culture. Such enrichment may be possible through FACS sorting pre- or post-heat killing to isolate spore-sized cells for time-lapse imaging during germination. In addition to differences in colony coloration during sporulation between the lab strain 168 and the biofilm-forming strain 3610, strain 3610 also sporulated more readily in LB (Fig. 4C,D). Thus, although sporulation has been extensively studied in laboratory strains such as 168, the biofilm-forming strain 3610 and other environmental isolates may provide the key to further mechanistic insights into this process.

Given the observation that knockdown of fatty acid synthesis caused hyperwrinkled colonies (Fig. 3C) and reduced sporulation efficiency (Fig. 4F,H), future investigations should focus on whether the regulation of fatty acid synthesis mechanistically connects wrinkling and sporulation. Regardless, these findings add to a growing body of evidence linking fatty acid synthesis to a wide range of cellular processes in diverse bacteria. Fatty acid synthesis dictates cell size in *Escherichia coli* (31, 32), impacts biofilm formation in *Salmonella* (33), and is nonessential in some pathogens when host fatty acids are available (34). Improved understanding of the links between the biochemical effects of modulating fatty acid levels and other consequences such as changes to membrane potential may help to interpret our systems-level phenotypic quantifications. Together, our results demonstrate that CRISPRi essential gene knockdowns are an effective tool to uncover the role these genes play during biofilm and non-biofilm growth and development of *B. subtilis*.

## Methods

### Media

Strains were grown in LB (Lennox broth with 10 g/L tryptone, 5 g/L NaCl, and 5 g/L yeast extract) or MSgg medium (5 mM potassium phosphate buffer, diluted from 0.5 M stock with 2.72 g K_2_HPO_4_ and 1.275 g KH_2_PO_4_, and brought to pH 7.0 in 50 mL; 100 mM MOPS buffer, pH 7.0, adjusted with NaOH; 2 mM MgCl_2_•6H_2_O; 700 μM CaCl_2_•2H_2_O; 100 μM FeCl_3_•6H_2_O; 50 μM MnCl_2_•4H_2_O; 1 μM ZnCl_2_; 2 μM thiamine HCl; 0.5% (v/v) glycerol; and 0.5% (w/v) monosodium glutamate). MSgg medium was made fresh from stocks the day of each experiment for liquid cultures, or a day before the experiment for agar plates. Glutamate and FeCl_3_ stocks were made fresh weekly. Colonies were grown on 1.5% agar plates. TY medium was made for phage transduction using the LB recipe above supplemented with 10 mM MgSO_4_ and 0.1 mM MnSO_4_. DSM medium was made with 8 g/L Difco nutrient broth with 0.5 mL/L 1 M MgSO_4_, 10 mL/L 10% KCl, and 0.5 mL/L 1 M NaOH. After autoclaving and cooling, 1 mL/L 1 M Ca(NO_3_)_2_, 1 mL/L 0.1 M MnCl_2_, and 1 mL/L 1 mM FeSO_4_ were added. Xylose at 1% final concentration was used for full knockdowns. Antibiotics were used where indicated at the following concentrations: ampicillin (amp, 100 µg/mL), MLS (a combination of erythromycin at 0.5 μg/mL and lincomycin at 12.5 μg/mL), chloramphenicol (cm, 5 μg/mL), tetracycline (tet, 12.5 μg/mL) and spectinomycin (spc, 100 μg/mL).

### Strain construction

All strains and their genotypes are listed in Table S1. The RFP-depletion strain (HA13) was constructed using phage transduction (35), using HA12 as a parent and HA11 (CAG74226) as the donor. A 168 strain containing *P_xyl_-dcas9* at the *lacA* locus (CAG74399) was used as a donor and wild-type strain 3610 was used as the recipient to create the 3610-dCas9 parent strain (CAG74331/HA2) using MLS for selection.

For CRISPRi library construction, the 3610-dCas9 parent strain was used as the recipient and strains from a 168 CRISPRi library (13) were used as the donor. The phage transduction protocol was amended to increase the throughput of strain construction as follows. Donor strains were grown in 96-well deep-well plates (1-mL cultures in TY medium) for at least 5 h with shaking at 37 °C and a Breathe-easy (Sigma-Aldrich) film covering the plate. 0.1 mL of 10^-5^ dilutions of fresh phage stocks grown on strain 3610 cells (10^-5^ was chosen as the dilution factor because it provided the appropriate level of lysis for our phage stock in a trial transduction) were aliquoted into 77 or 71 glass test tubes (each plate of the library contains 77 strains, except the fourth plate that contains 71 strains). 0.2 mL of each culture were added to the tubes and the entire rack was incubated at 37 °C for 15 min. Then, working quickly in batches of 11, 4 mL of TY molten soft agar (∼55 °C) were added to each phage-cells mixture, mixed gently, and poured onto TY plates so that the soft agar covered the entire plate. These plates were incubated at 37 °C overnight in a single layer (not stacked). The next day, the plates were examined for lysis and 5 mL TY broth with 250 ng DNase were added to each plate and the top agar was scraped with a 1-mL filter tip to liberate phage. The TY broth was then pipetted into a syringe attached to a 0.45-µm filter and carefully filtered into a 5-mL conical vial. After filtering, 1 mL of lysate was added to the appropriate well of a 96-well deep-well plate. Once all of the phage was isolated, we arrayed 10 µL of each phage stock into 96-well microtiter plates. One hundred microliters of a saturated (>5 h of culturing, OD_600_>1.5) culture were aliquoted into the wells containing phage and incubated for 25 min at 37 °C without shaking. The phage/cell mixtures were plated onto selection plates (LB with chloramphenicol and citrate to select for the sgRNA locus), which were incubated for 18 h at 37 °C. Any plates that did not have visible colonies after this incubation were incubated further at room temperature, and colonies generally appeared within a day. Transductant colonies were streaked for single colonies on LB+chloramphenicol plates and a single colony for each strain in the library was stocked by growing in 5 mL LB on a roller drum at 37 °C to mid-to late-log phase and then adding the culture to the appropriate well of 96-well plate with a final concentration of 15% glycerol. The library was stored at -80 °C.

To generate an *mNeongreen* reporter plasmid pKRH97, the promoter region of *hag* was PCR-amplified from DK1042 chromosomal DNA using primers 5838 (aggagggatccttatcgcggaaaataaacgaagc)/5839 (ctcctctcgaggaatatgttgttaaggcacgtc) and digested with BamHI and XhoI. Next, the *mNeongreen* gene was PCR-amplified from DK3394 using primers 5836 (aggagctcgagtaaggaggattttagaatggtttcgaaaggagaggag)/5837 (ctcctgaattcttacttatagagttcatccatac) and digested with XhoI and EcoRI. Both products were simultaneously ligated into the BamHI and EcoRI sites of pKM086 containing a polylinker and tetracycline resistance cassette between two arms of the *ycgO* gene (36). To generate the *P_eps_-mNeongreen* reporter plasmid pDP565, the promoter region of *epsA* (*P_eps_*) was PCR-amplified from DK1042 chromosomal DNA using primers 7524 (aggagaagcttcctgtcgttatttcgttcatta)/7525 (ctcctctcgagccctttctgttaatgattggatt). The PCR product was digested with HindIII and XhoI and cloned into the HindIII and XhoI sites of plasmid pKRH97. To generate the *P_tapA_-mNeongreen* reporter plasmid pDP575, the promoter region of *tapA* (*P_tapA_*) was PCR-amplified from DK1042 chromosomal DNA using primers 7612 (aggagaagcttccttttggctataaggatcaaatg)/7613 (ctcctctcgaggtaaaacactgtaacttgatatg). The PCR product was digested with HindIII and XhoI and cloned into the HindIII and XhoI sites of plasmid pKRH97. Strain DK1042 (37) was transformed with the above plasmids to integrate the constructs into the genome. The *Peps-mNeongreen* construct was then transduced into DS6988, a Δ*sinR* Δ*epsH* recipient strain.

The amended phage transduction protocol above was used to make the CRISPRi candidate gene depletion reporter strains and the sporulation strains. Phage transduction was used to introduce the *P_epsH_* (HA1463 donor) and *P_tapA_* (HA1502 donor) reporter constructs into parent strain HA2 containing *P_xyl_-dcas9* to create parent strains HA1464 (parent *P_epsH_*) and HA1505 (parent *P_tapA_*). These parents were used as recipient strains and the strain 168 CRISPRi strains as donors for phage transduction (13). HA395 and HA396 were used as parents to introduce the reporter constructs into Δ*sinI* and Δ*sinR* strains, respectively. For sporulation strains, we used strain 3610 (HA10) as the recipient and strains HA1163-HA1172 as donors for phage transduction (Table S1).

### CRISPRi targeting of RFP in colonies and biofilms

Wild-type 3610, parent-RFP, and CRISPRi-RFP strains were cultured in 5-mL test tubes at 37 °C to an OD_600_∼1 in liquid LB. The parent-RFP strain was spotted onto LB and MSgg agar plates without xylose, while the CRISPRi-RFP strain was spotted onto LB and MSgg agar in 12-well plates containing 0.0005% to 1% xylose. Each well of the plate had ∼1 mL media to maintain the proper focal plane. Colonies were incubated at 30 °C for 48 h inside a box in a warm room to prevent the plates from drying out. RFP fluorescence of the colonies was imaged with a Typhoon FLA 9500 scanner using the multi-plate drawer, with plates inserted face up so the scanner imaged through the agar. RFP signal was acquired with a 532-nm laser and a long-pass green filter. FIJI was used to quantify the fluorescence intensity of each colony: the images were first inverted so the background would be black and RFP signal would be white, and fluorescence was measured using a circle the size of the largest colony. The wild-type 3610 measurement was subtracted as a blank.

### Liquid growth assays for the CRISPRi library

Freezer stocks were pinned onto agar plates using sterile 96 long-pin Singer RePads (REP-001) and grown overnight at 37 °C to form colonies. Sterile 96 long-pin Singer RePads were used to manually inoculate the colonies into 200 µL of LB in a 96-well microtiter plate.

Parent controls were added to empty wells on each plate. The plate was sealed with optical film and a syringe needle was used to poke off-center holes above each well to allow air exchange in the headspace and grown with linear shaking at 37 °C for ∼5 h until the parent control reached an OD_600_∼1. The plate was diluted 1:200 into 200 µL of the appropriate medium and growth was monitored on a Biotek Epoch plate reader at 30 °C with linear shaking (567 cycles per min, 3-mm magnitude). The minimum value of a blank well was used as the blank.

### Liquid growth assays for follow-up experiments

Strains were streaked out for individual colonies from glycerol stocks onto LB-agar plates and incubated at 37 °C overnight. Single colonies were outgrown in 200 µL of LB in a 96-well microtiter plate at 37 °C for ∼5 h until WT/parent strains reached an OD600∼1. Cultures were diluted 1:200 into 200 µL of the appropriate medium, the plate was sealed with an optical seal with holes poked off-center for air exchange, and growth was monitored using a Biotech Epoch plate reader at 30 °C (*asnB* knockdown) or 37 °C (*mapA*, *glyA*, *folD* knockdowns) with linear shaking (567 cycles per min, 3-mm magnitude). Before diluting, the medium+microtiter plates were pre-blanked to enable accurate measurements of growth (38).

### Growth assay of colonies on agar plates

Freezer stocks were pinned onto agar plates using sterile 96 long-pin Singer RePads and grown overnight at 37 °C to form colonies. Sterile 96 long-pin Singer RePads were used to inoculate the colonies into 200 µL of LB in a 96-well microtiter plate. Parent controls were added to empty wells on each plate. The plate was sealed with an optical seal with holes poked off-center for air exchange and grown with linear shaking at 37 °C for ∼5 h until the parent control reached an OD_600_∼1. Approximately 1 µL was spotted onto an agar plate (poured the day before, dried at room temperature overnight, and then dried at 37 °C for 1 h before spotting) using a Singer robot (spotting was repeated 12 times with a Singer RePad) and colonies were grown at 30 °C in a single layer inside a box or plastic bag to prevent the plates from drying out. Images were taken 24 h post-spotting for colony size analyses and 48 h post-spotting for biofilm wrinkling analyses. For follow-up assays of candidate gene knockdown strains, the same protocol was used except that freezer stocks were streaked out for individual colonies that were used for inoculation.

### Colony imaging

Images were acquired with a Canon EOS Rebel T5i EF-S with a Canon Ef-S 60 mm f/2.8 Macro USM fixed lens. The DSLR camera was set up at a fixed height in a light box. Lighting and camera settings were maintained for the duration of the experiment, using the “manual” mode on the camera. The EOS Utility software was used to run the camera. Plates were imaged colony side up to avoid imaging through the agar.

### Colony size analysis

Colony size was measured in FIJI by diagonally bisecting the center of the colony. Colony size was assayed based on images acquired at 24 h, which was before the colonies were noticeably affected by the growth of neighbors or the edge of the plate.

### Wrinkling quantification

Images were analyzed with custom Matlab code (https://doi.org/10.25740/nt084hv1801). In brief, images were rotated so that the boundaries of the plate were aligned with the horizontal and vertical axes. Images were converted to grayscale. To account for uneven lighting across the image, the background intensity profile was computed using a disk-shaped structured element of size 15 pixels, and was subtracted from the image.

Background-subtracted images were contrasted-adjusted and converted to a binarized image using `im2bw` with default parameters. These binarized images were typically a reasonable proxy for the positions of wrinkles. All connected components with fewer than 50 pixels were removed using `bwareaopen`. The center of one of the corner colonies was manually recorded using `ginput`, and the centers of the other colonies were inferred based on the known grid spacing. The wrinkling intensity was computed as a function of radial distance *r* by summing the pixels within a circle of radius *r*. The value used to calculate the reported wrinkling density was the radius of the colony as measured in FIJI, and the total intensity was normalized to colony area. Note that the *sufD* strain was mis-classified due to a lighting artifact (Fig. S3A).

### mNeongreen reporter assay

Strains from glycerol stocks were streaked out for single colonies and plates were incubated at 37 °C overnight. Individual colonies were picked into 200 µL of LB medium in a 96-well plate and grown at 37 °C with linear shaking until the parent/wild-type controls reached an OD_600_ ∼1. One microliter of culture was spotted onto MSgg agar rectangular plates (poured the day before, dried overnight at room temperature and 1 h at 37 °C before spotting; ∼30 mL of medium were used). Plates were incubated in a box at 30 °C for 16 h.

Plates were imaged with a Typhoon FLA 9500 scanner using the multi-plate drawer, inserted face up so that the scanner imaged through the agar. mNeongreen signal was acquired with a 473-nm laser and a long-pass blue filter. Images were inverted so the background was dark and fluorescence was in white, and fluorescence intensity was measured in FIJI using a circle the size of the largest colony. Colony size was measured diagonally with a line bisecting the center of the colony to calculate intensity per unit area.

### Essential gene bioinformatic comparison

A bash script was used to compare the protein sequences of essential genes in strain 168 (PRJNA57675_168.fasta) with their homologs in strain 3610 (PRJNA377766_3610.fasta), and any mismatches were identified using custom Matlab code. All scripts are available in the repository https://purl.stanford.edu/fb108cz5121. Any genes with potential protein coding differences between strains 168 and 3610 were tested using BLAST (https://blast.ncbi.nlm.nih.gov, NCIB strain 3610 genome GenBank: CP034484.1, strain 168 genome from SubtiWiki (39)).

### Sporulation assay

The library was grown from colonies at 37 °C for ∼5 h so that the wild-type strain reached an OD_600_∼1 in a plate reader with linear shaking. Cultures were diluted 1:200 either using a Singer robot (for DSM experiments) or by manually pipetting (LB) into DSM or LB, respectively, and grown for 24 h at 37 °C in a Biotek Epoch plate reader with linear shaking (567 cycles per min, 3-mm magnitude). At 24 h, vegetative cells were heat-killed by transferring the sealed plates to an oven at 80 °C for 30 min with pre-heated heat blocks placed on top of the plates to prevent condensation on the seal. Seals were carefully removed post-heat-kill and cultures were diluted 1:200 into LB. To monitor germination, cultures were grown at 37 °C in an Epoch plate reader with linear shaking for 24 h. Plates were pre-blanked before inoculation to enable accurate measurements of growth (38). For follow-up assays to confirm delayed growth phenotypes, strains were streaked out for individual colonies, plates were incubated overnight at 37 °C, and individual colonies were inoculated into each well of a 96-well plate for outgrowth following the sporulation assay as described above.

### Sporulation and plating efficiency

Cultures were grown as above in the sporulation efficiency assay. Serial dilutions of cultures before and after heat-killing were plated to quantify total viable cells and surviving spores, respectively. Seals were removed to perform the serial dilution, then plates were resealed with a new seal before heat-killing. Cultures were diluted with 10 µL into 90 µL in v-bottom plates and pipetted up and down several times to mix using new pipette tips for each dilution. Ten microliters of each dilution were spotted using a BenchSmart (Rainin) liquid handling robot. Plates were incubated overnight at 37 °C, and colonies were counted. For the spore dilution experiment, plating efficiency of undiluted heat-killed spores was measured as described above. Ten-fold serial dilutions of the heat-killed culture were then used to inoculate (1 µL into 200 µL) LB and growth was monitored as described above.

### Determining genes required for growth in liquid and agar

Twelve strains were unable to grow under full induction in liquid despite being able to grow as a colony on LB+xylose and MSgg+xylose. However, these gene knockdowns exhibited 10^3^- to 10^4^-fold reductions in plating efficiency (performed as described above with serial dilutions and plating) relative to the parent strain (Fig. S2D,E), consistent with their growth defects in liquid. Thus, these 12 strains were presumed to be incorrectly classified based on our colony screen.

## Acknowledgements

The authors thank the Huang lab and Zachary Hallberg for helpful discussions. The authors acknowledge support from the Allen Discovery Center at Stanford on Systems Modeling of Infection (to H.A.A. and K.C.H.), NIH R35 GM118061 (to C.A.G.), NIH RM1 Award GM135102 (to K.C.H.), and NSF Awards EF-2125383 and IOS-2032985 (to K.C.H.). K.C.H. is a Chan Zuckerberg Biohub Investigator.

## Supplementary Figures

**Figure S1:**
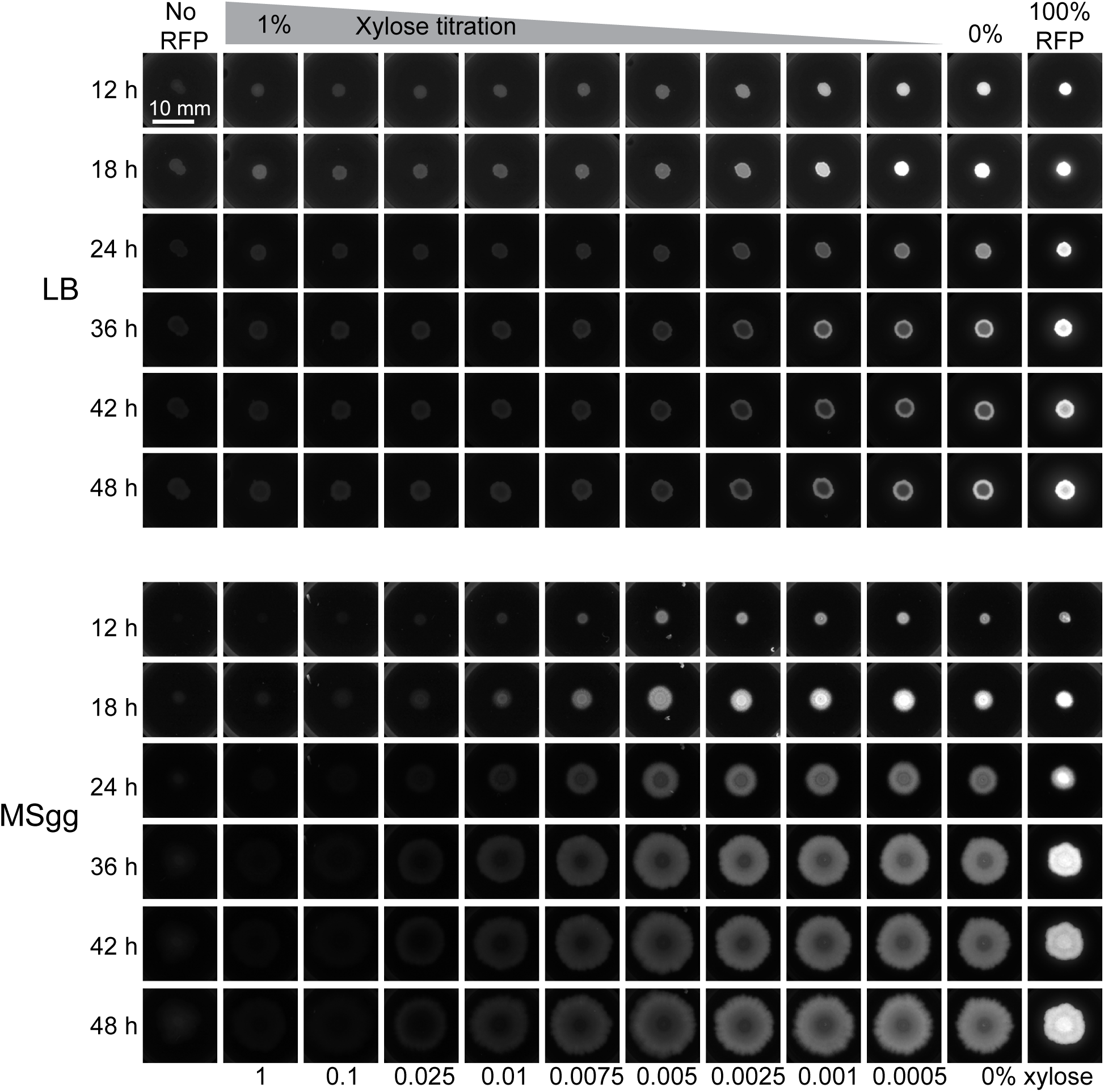
CRISPRi effectively inhibits RFP expression over 48 h in colonies. Images of RFP fluorescence over time in colonies growth on LB and MSgg with various concentrations of xylose. These colonies were used to generate the data in Fig. 1C.

**Figure S2:**
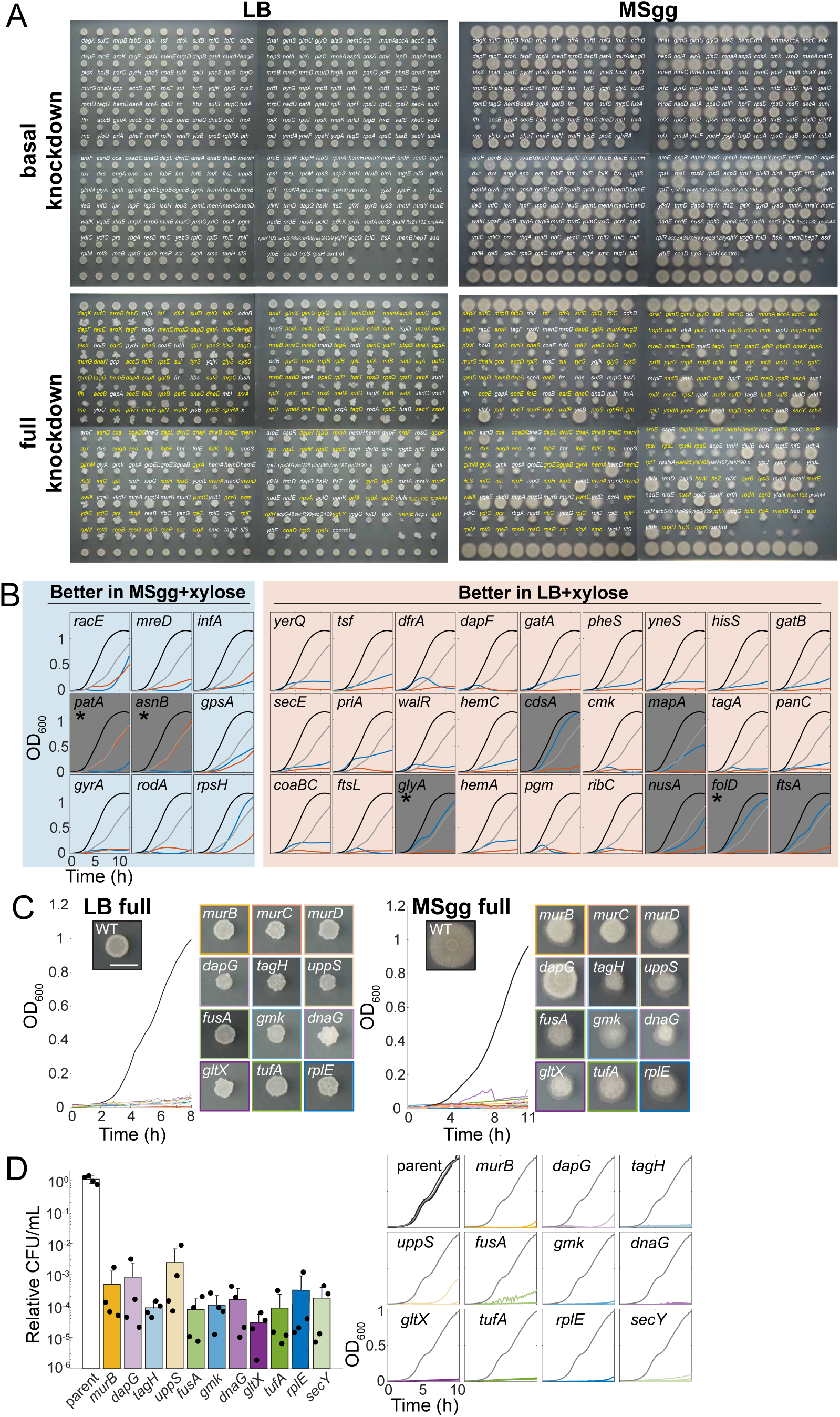
Growth phenotypes of knockdown strains on surfaces and in liquid. A) All strains grow on agar plates in basal knockdown conditions (top), but many are inhibited for growth in full knockdown conditions (+1% xylose, bottom). Controls were spotted on the top and/or bottom of each plate (top and bottom rows of the combined images). Images were taken at 24 h. Full knockdown colonies were scored as not growing (dying, not growing outside of the original spot, or generating suppressors) or growing outside of the original spot. Genes scored as not growing as colonies are annotated on the full depletion plates in yellow, and non-essential genes that exhibited poor growth are labelled with an asterisk. B) Most strains that exhibited growth in one medium (LB or MSgg) grew poorly in the other medium. Data shown are strains that grew above the OD_600_ threshold in one medium during the initial screen. However, for most strains growth was only slightly above the threshold; gray boxes denote the strains with substantial growth differences between the two media. Of these, only the strains marked with asterisks had a reproducible phenotype and hence were considered as true positives (Fig. 2B). Wild-type growth curves in LB+1% xylose or MSgg+1% xylose are shown in black or gray, respectively. Knockdown growth curves in LB+1% xylose or MSgg+1% xylose are shown in blue or red, respectively. C) Strains that appeared to grow as colonies (right) but not in liquid (left) from the original screen. Scale bar: 5 mm. Wild-type curves are shown in black and the mutants are shown in various colors. D) Strains that appear to grow as colonies but not in liquid are reduced for colony forming ability (left). Each strain was grown in LB and plated on LB+1% xylose plates and diluted into LB+1% xylose (right). Mutant growth curves are show in various colors as in (C) while the average of the parent growth curves is shown with one black line, except for the ‘parent’ panel in which all four growth curves are shown. *n*=4 biological replicates.

**Figure S3:**
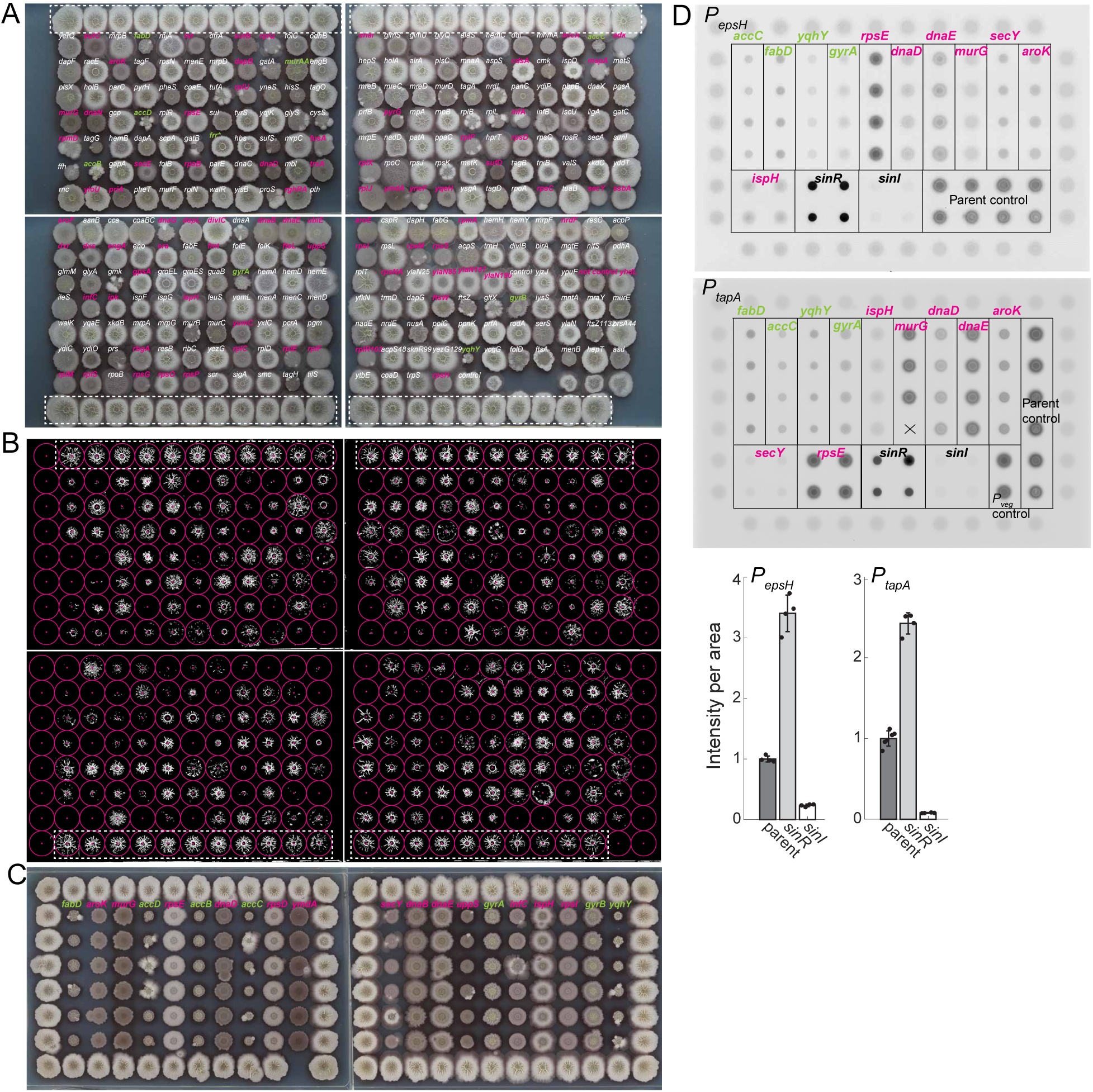
Knockdown strains display various wrinkling phenotypes. A) Biofilm colonies exhibit a wide range of wrinkling. Cultures were spotted on MSgg agar and grown for 48 h to form biofilm colonies. Controls are outlined in white dashed boxes. High wrinkling strains are labeled with green text and low wrinklers are labeled in pink. *sufD* was identified by eye but through image analysis, likely due to contrast. B) Images in (A) after application of our analysis pipeline (Methods) that highlights wrinkles as white pixels. Colony wrinkling was quantified based on the total number of white pixels inside each circle surrounding a colony. Controls are outlined in white dashed boxes. C) The phenotypes of select strains with high and low wrinkling were validated. Wild-type controls were spotted in the exterior wells. *n*=6 biological replicates for each mutant. D) Expression from matrix reporters shows that matrix expression is generally uncorrelated with wrinkling. CRISPRi mutants expressing mNeon expressed from the *epsH* (top) or *tapA* (middle) promoters were spotted onto MSgg agar and colonies were imaged at 16 h. Bottom: intensity per unit area for the parent and positive (Δ*sinR*) and negative (Δ*sinI*) controls. Unlabeled wild-type cells were spotted in the outside wells and at least three biological replicates of each mutant were included.

**Figure S4:**
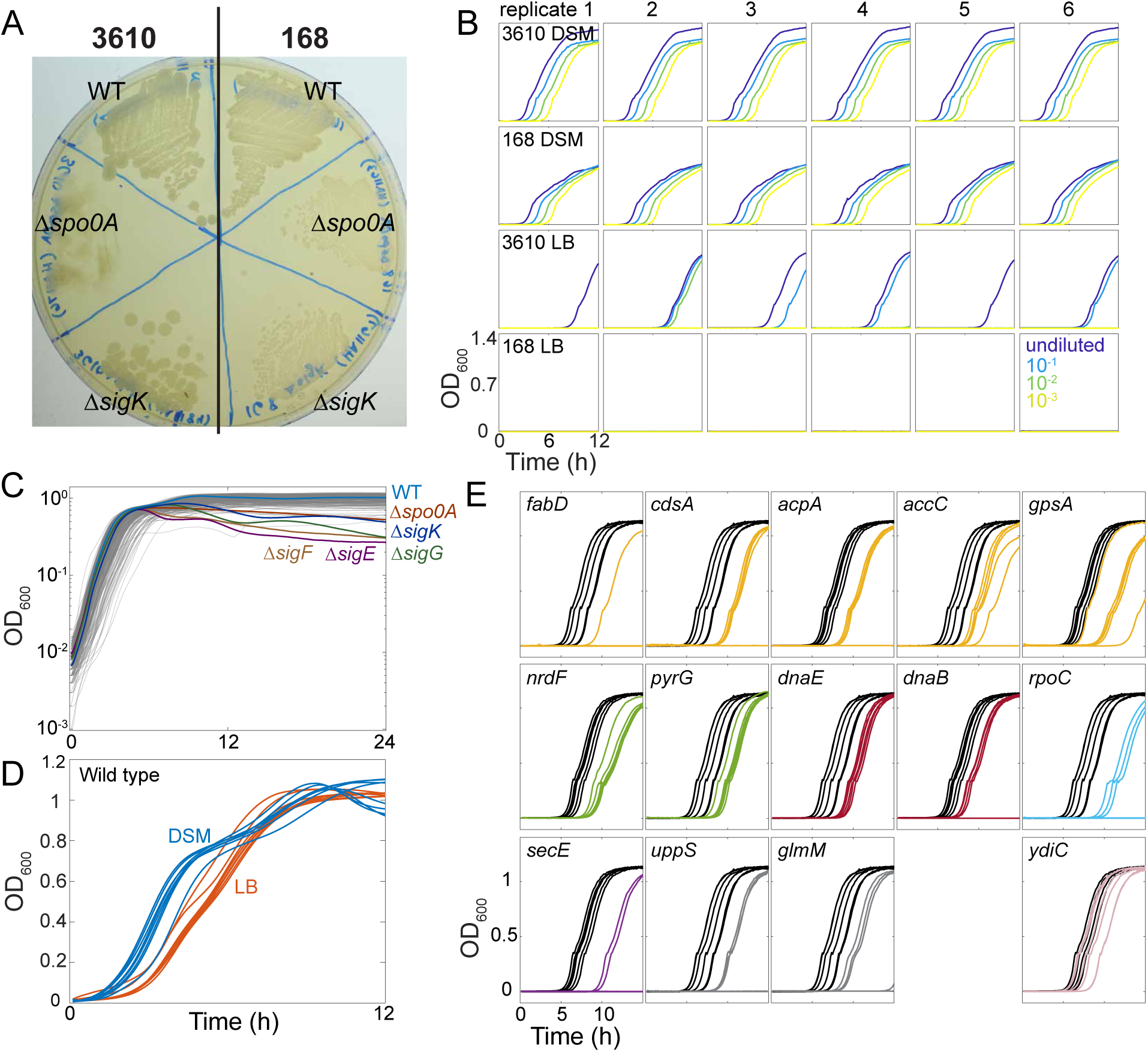
A high-throughput sporulation screen reveals gene depletions that reduce sporulation in LB. A) Strain 3610 sporulation mutants are opaque while strain 168 mutants are more transparent than wild type. Strain 3610 and strain 168 wild type, Δ*spo0A,* and Δ*sigK* were struck onto DSM agar plates and incubated overnight and then imaged using a lightbox. B) Growth curves of post-heat-kill cultures demonstrate that the delay in outgrowth is correlated with the inoculum size. Strains were grown prior to heat-killing in DSM or LB media. The strain 168 LB cultures were below the limit of detection for our assay. C) None of the knockdown mutant growth curves resemble those of known sporulation mutants in DSM. Growth curves of the known mutants are shown as bold colored lines, while library strains are shown in gray. D) Wild-type strain 3610 cultures exhibit distinct growth curves in DSM (blue) and LB (red). Six biological replicates are shown. E) Reduced sporulation/germination phenotypes of knockdowns are reproducible. Shown is growth of six biological replicates of each knockdown (yellow) and wild type (black). No growth indicates that that sporulation levels were below the limit of detection of our assay. Δ*ydiC* was included as a positive control for sporulation, and its curves overlapped with those of wild type. Fatty acid mutants are colored in yellow, DNA replication-related mutants are red, ribosome-related genes are blue, cell wall-related genes are gray, biosynthesis mutants are green, and a secretion mutant is purple.

